# Analysis of Maize Profilin-4 Isoform as an allergen

**DOI:** 10.1101/503425

**Authors:** Saruar Alam, Md. Kamrul Hasan, Md. Faruk Hossain

**Affiliations:** Department of Biochemistry and Molecular Biology, University of Dhaka, Dhaka, Bangladesh; Department of molecular neuroscience, St. John’s University, Queens, New York 11439

**Author notes:** Equal contribution (Saruar Alam, Md. Kamrul Hasan).

**Keywords:** Profilin-4, allergenicity, *Zea mays*, allergen, in silico, epitope

## Abstract

Profilin is an actin monomer-binding protein that controls the dynamic turnover of actin filaments and is ubiquitously present in different organisms ranging from prokaryotes to higher eukaryotes. Maize (Zea mays) profilin-4 isoform is a pollen-specific protein. Birch profilin isoform is a known allergen but maize profilin is yet to be characterized. Due to the high cultivation rate of maize, the analysis of the properties of maize profilin-4 isoform as an allergen is a demand of time for developing an effective immune therapy. Here, we analyzed the allergenic potency of profilin-4 by studying of its physicochemical properties, including molecular weight (∼14kD) and theoretical pI (4.63). We tested the potential B cell epitope candidates of profilin-4 using different immune-informatics tools housed at IEDB analysis resource. For the B cell epitope prediction, potential antigenic sites on the protein surface were predicted by both propensity scale and machine learning method followed by their mapping of 3D structure prediction. We hereby claim that the profilin-4 is a potential allergen and is able to induce allergic responses. However, the wet lab experiments are needed to validate our claim that’s beyond our scope.

## 1. Background

Allergens are small proteins or glycoproteins wavering a molecular weight range of 15 to 40 kDa [1]. Allergens appear from different sources, for instance, pollen allergens from plants, venom allergens from insects, food allergens from various food items, mite allergens from dust, etc. [2]. They can induce IgA, IgE, IgG, and IgM antibody-mediated immune responses [3]. In addition, they can induce Th2 (Helper T) cell-mediated immune response in the human body [4]. Allergens produce an enzymatic or immunogenic reaction to cause allergenicity [5].

*Zea mays* (maize), a Poaceae family member, is one of the most cultivated crop plants around the world. Maize has both nutritional and medicinal importance. The maize kernel is the nutritive part of the plant that contains all the different vitamins, fatty acids, minerals, etc. Maize is a great source of phytochemicals that are used to treat choric diseases, HIV, even cancer, etc. [29]. There is an increasing trend of maize production over the last decade. It has been estimated that about 187.95 million hectors of land were used for maize cultivation[6].

In this study, we predict profilin-4 as a potential allergen. As the cultivation rate of maize increases keeping the pace with the demands, it also provokes the concern of allergenicity of its pollen. Wind-pollinated seed plants produce pollens which encompass crucial sign of Type-I allergy [7]. Profilin is known as panallergen due to its widespread cross-reactivity [11]. The allergenic properties of pollens have no association with biological function but the enzymatic and immunogenic actions of allergens cause the allergic reaction and inflammation [8]. The profilin of birch pollen (called Bet v 2) [10] and latex [9] are documented as allergens, but not the maize-specific profilin isoform, profilin-4. Using the Bioinformatics tools and database [12–16], here we analyzed the allergenicity of profilin-4.

## 2. Methodology

### 2.1 Protein Sequence retrieval

Profilin-4 protein sequence (O22655.1) for *Zea mays* was retrieved in FASTA format from NCBI protein database (http://www.ncbi.nlm.nih.gov/protein). This protein sequence was further used to perform different computational analysis from linear amino acid residues.

### 2.2 Prediction of physicochemical properties

Different physicochemical properties for profilin-4 protein was predicted from its linear amino acid sequence using ProtParam tool (https://web.expasy.org/protparam/) web server. Protparam predicts the molecular weight, theoretical pI, atomic composition, amino acid composition, instability index, extinction coefficient, grand average of hydropathicity (GRAVY), estimated half-life, and aliphatic index of any given amino acid sequences [17].

### 2.3 Potential antigenic sites prediction

The hydrophobic and hydrophilic regions were determined to predict the antigenicity of profilin-4. The hydrophilic portions are exposed to the surface of the protein and display reactivity to the B cell. Kolaskar-Tongaonkar antigenicity and Parker’s hydrophilicity were used to predict the antigenicity of profilin-4. Antigenic propensity, as well as hydrophilicity, was then analyzed from the plots generated. [18, 19].

### 2.4 Potential B cell epitope prediction

Not all the regions exposed to outer surface react with B cell, that’s why to predict the B cell epitopes a machine learning tool was used (http://tools.iedb.org/bcell/), a web server where Bepipred linear epitope prediction method was chosen [20]. Bepipred linear epitope prediction method uses an algorithm comprising both hidden Markov model and antigenic propensity and thus allowed to cross check the predicted result from Kolaskar-Tongaonkar antigenicity and Parker’s hydrophilicity prediction method [18,19].

### 2.5 Prediction of 3D structure of profilin-4 and mapping of B cell epitopes on the predicted structure

For 3-D structure prediction of profilin-4 an online tool RaptorX (http://raptorx.uchicago.edu) web server was used. RaptorX does several alignments of the target protein sequence with different protein templates to predict a model [21-22]. We found that the structure (PDB ID: O22655) of profilin-4 showed maximum alignment score with its target template. Swiss PDB tool was used for the energy minimization of the structure [23]. To validate the structure a Ramachandran plot was generated using an online tool RAMPAGE (http://mordred.bioc.cam.ac.uk/~rapper/rampage.php) web server which measures the stereo-chemical properties of the protein structure [24]

## 3. Results and Discussion

### 3.1 Physicochemical properties predict allergenic property of profilin-4 protein

Sometimes physicochemical properties of a protein can determine the allergenic property of a protein [25]. The maize profilin-4 consists of 131 amino acids with molecular weight of approximately 14 kD (Table 1). The total amino acid distribution of profilin-4 protein (Figure1) shows that asparagine present in the lowest amount and glutamate, glycine, isoleucine, and valine predominate among the 20 amino acids of profilin-4 protein (Figure 1). Due to abundant acidic amino acids, this suggests the protein’s theoretical pI is to be acidic and theoretical pI found 4.63 which means profilin-4 protein is highly acidic and tends to be allergenic [25]. Hence negatively charged residues (Asp + Glue) is twice the total number of positively charged residues (Arg + Lys) (Table 1) in profilin-4, there is probability to be processed by dendritic cells via scavenger receptor [26]. From predicted half-life and instability index it indicates that profilin-4 is quite stable [27]. From the predicted negative grand average of hydropathicity value, it can be assumed that most of the amino acid residues of profilin-4 protein are likely to be present on the surface of the folded profilin-4.

**Table 1:** ProtParam predicted physicochemical properties for profilin-4 protein from *Zea mays*

| Properties | score |
| --- | --- |
| Number of amino acids | 131 |
| Molecular weight | 14101.15 |
| Theoretical pI | 4.63 |
| Total number of negatively charged residues<br>(Asp + Glu) | 18 |
| Total number of positively charged residues<br>(Arg + Lys) | 9 |
| Total number of atoms | 1971 |
| Ext. coefficient ($M^{-1}cm^{-1}$) | 17085 |
| The estimated half-life (hours) | 30 |
| The instability index (II) | 35.50 |
| Aliphatic index | 86.26 |
| Grand average of hydropathicity (GRAVY) | -0.020 |

**Figure 1:**
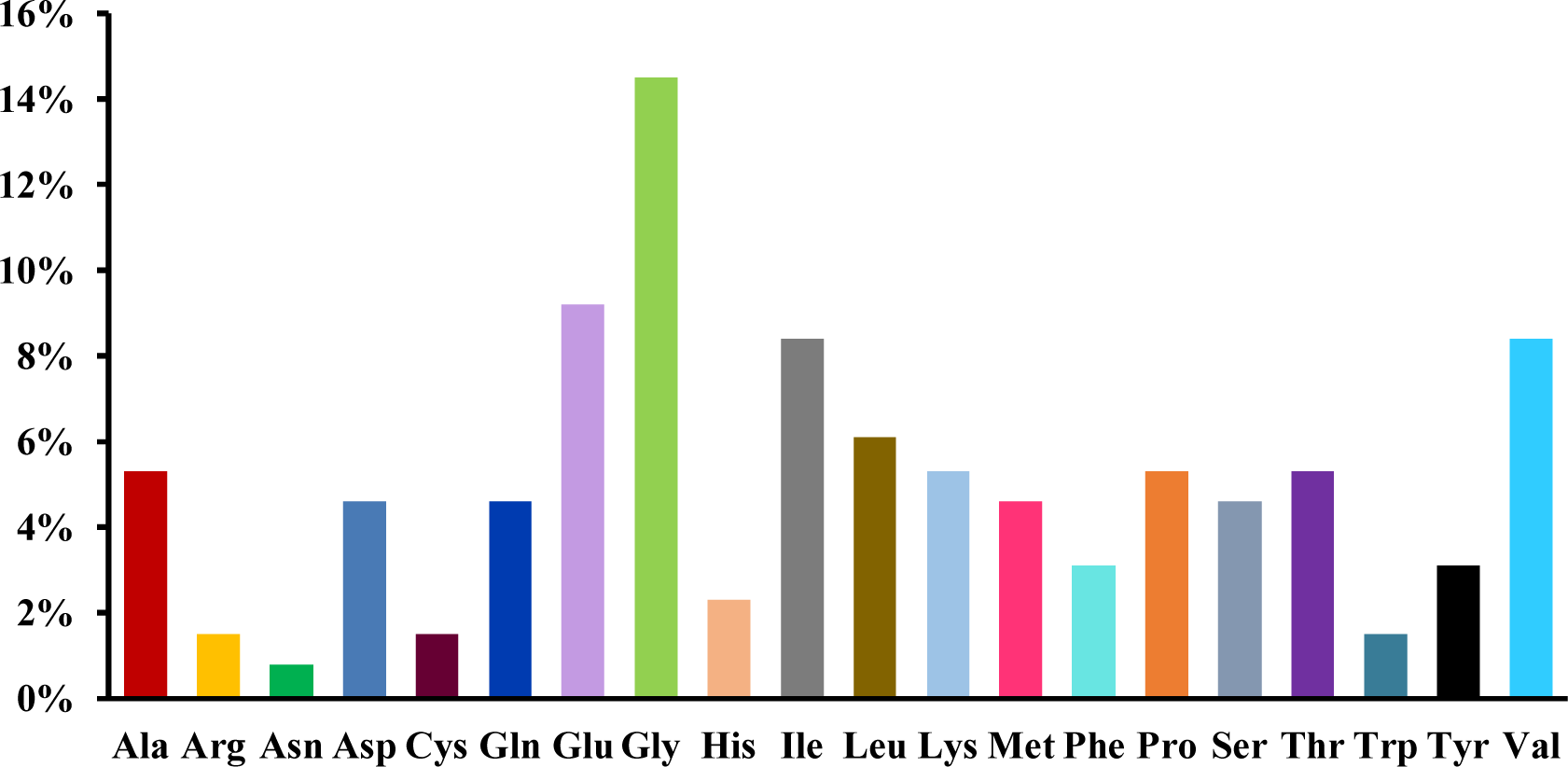
The amino acids composition in profilin-4. Glycine (Gly) and Asparagine (Asn) are the major (14.5%) and the least (∼0.8%) constituents, respectively

### 3.2 Prediction of Potential antigenic sites on the surface of the profilin-4 protein

For the prediction of profilin-4 allergenicity, Kolaskar and Tongaonkar prediction method were employed which functions on the basis of physicochemical properties of amino acids in proteins and abundances in experimentally known epitopes [18]. In Figure 2, the x-axis represents the amino acid position and the y-axis represents the antigenic propensity of the protein. The average antigenic propensity of profilin-4 protein is found to be 1.027. So all residues having a value greater than 1.027 are potential antigenic determinant. Seven peptides (Table 2) are found to be a potential antigen because they satisfy the set threshold value (1.00). The peptide regions “EGQHLSAAAIVGHDGSVWAQ” ranging from 16 to 35 amino acid residues and 100 to 108 amino acid residues (“SLIIGVYDE”) are predicted to have the highest antigenic propensity score. Both of them comprise about more than one-fifth (22.13%) of profilin-4 protein. The hydrophilic portion of a protein has a tendency to be exposed on the outer surface of the protein that makes them vulnerable to be engaged with B cell. The average score of hydrophilicity of profilin-4 is found to be 1.421 (Figure 3). The regions highlighted yellow have a hydrophilicity score of above the average and are likely to be present on the surface of the profilin-4 protein, while the regions highlighted green have a hydrophilicity score of below the average and are unlikely to be exposed on the surface. To predict the hydrophilic regions of profilin-4, we adopted Parker hydrophilicity prediction method [19]. For making a better prediction decision, we have also used a more reliable machine learning tool that follows the Bepipred linear epitope prediction method [28].

**Table 2:** The list of the Peptide sequences having at least 1.0 antigenic propensity score, predicted from Kolaskar and Togaonkar antigenicity plot

| Number | Start position | End position | Peptide sequence | Length |
| --- | --- | --- | --- | --- |
| 1 | 4 | 14 | QAYVDEHLMCE | 11 |
| 2 | 16 | 35 | EGQHLSAAAIVGHDGSVWAQ | 20 |
| 3 | 44 | 51 | PEEVAGII | 8 |
| 4 | 62 | 69 | PTGLFVGG | 8 |
| 5 | 72 | 85 | YMVIQGEPGVVIRG | 14 |
| 6 | 100 | 108 | SLIIGVYDE | 9 |
| 7 | 115 | 128 | CNMVVERLGDYLIE | 14 |

**Figure 2:**
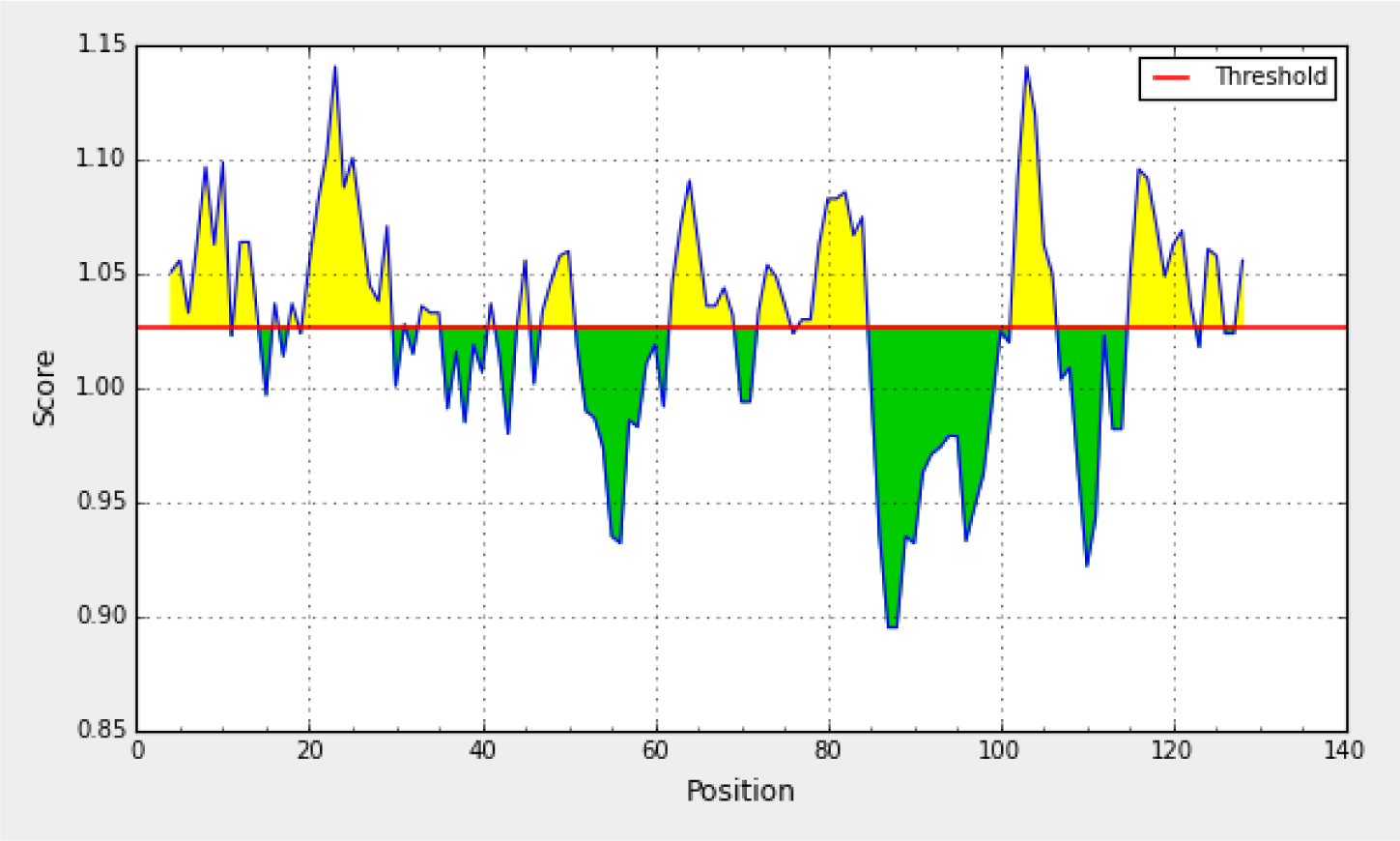
Kolaskar and Tongaonkar antigenicity graphical plot. The protein sequences that satisfied the set antigenic propensity threshold value of 1.00 are predicted to be potential antigenic region.

**Figure 3:**
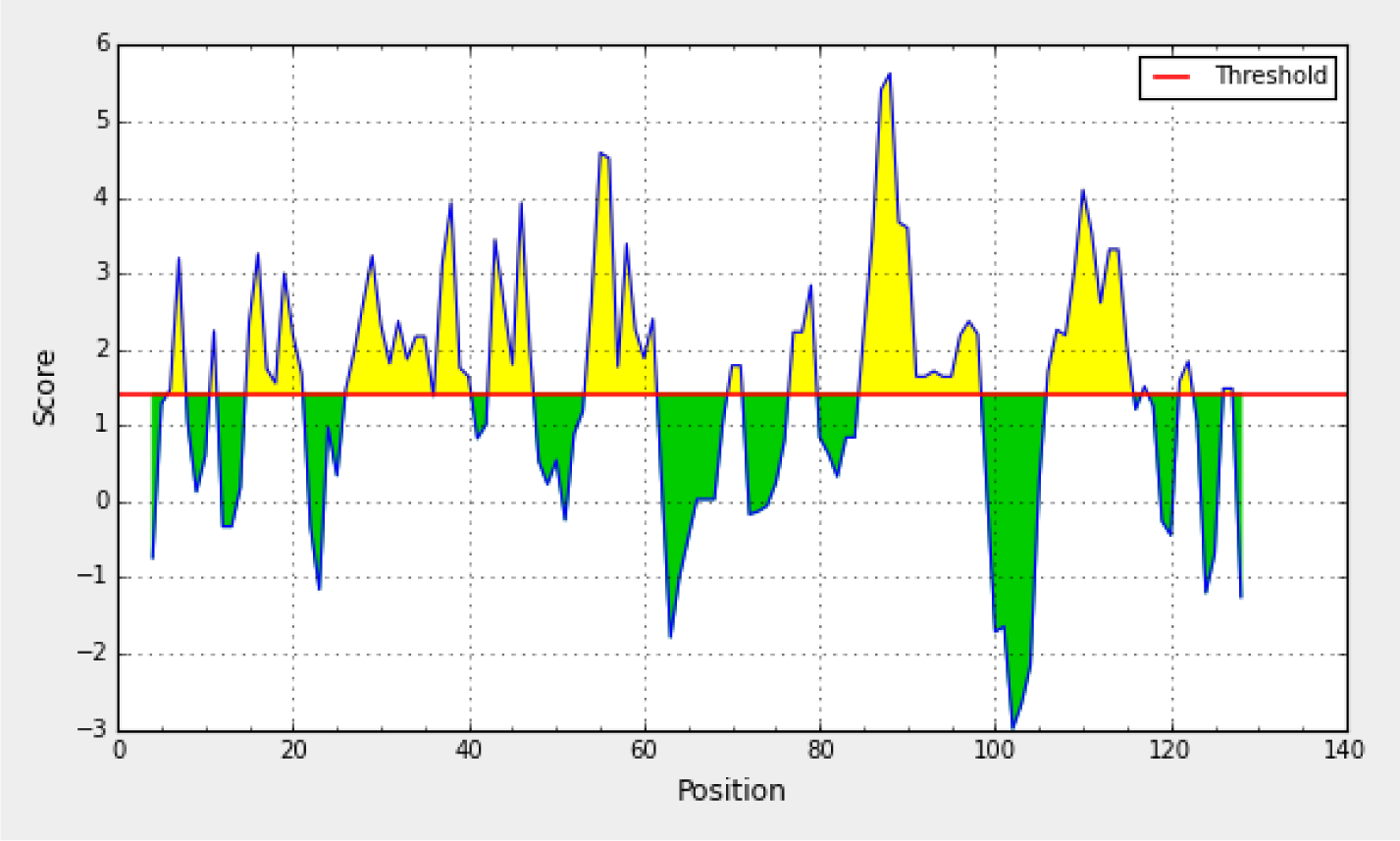
Parker hydrophilicity plot. The x-axis represents the amino acid position and the y-axis represents the hydrophilicity score.

### 3.3 Potential B cell epitopes overlap the antigenic sites of profilin-4

We have applied the BepiPred tool to predict the potential B cell epitopes. The Bepipred linear epitope prediction method uses an algorithm that links the Hidden Markov Model (HMM) and the antigenic propensity to make the prediction more trustworthy [20]. BepiPred predicted four potential B cell epitopes highlighted in yellow for profilin-4 protein sequence (Figure 4) and the maximum predicted score is 1.630. Predicted epitopes are summarized in Table 3.

**Table 3:** Predicted B cell epitope sequences and their position along with their length

| Number | Start position | End position | Epitope sequence | Length |
| --- | --- | --- | --- | --- |
| 1 | 30 | 47 | GSVWAQSESFPELKPEEV | 18 |
| 2 | 53 | 61 | DFDEPGTLA | 9 |
| 3 | 80 | 94 | GVVIRGKKGTGGITI | 15 |
| 4 | 107 | 114 | DEPMTPGQ | 8 |

**Figure 4.**
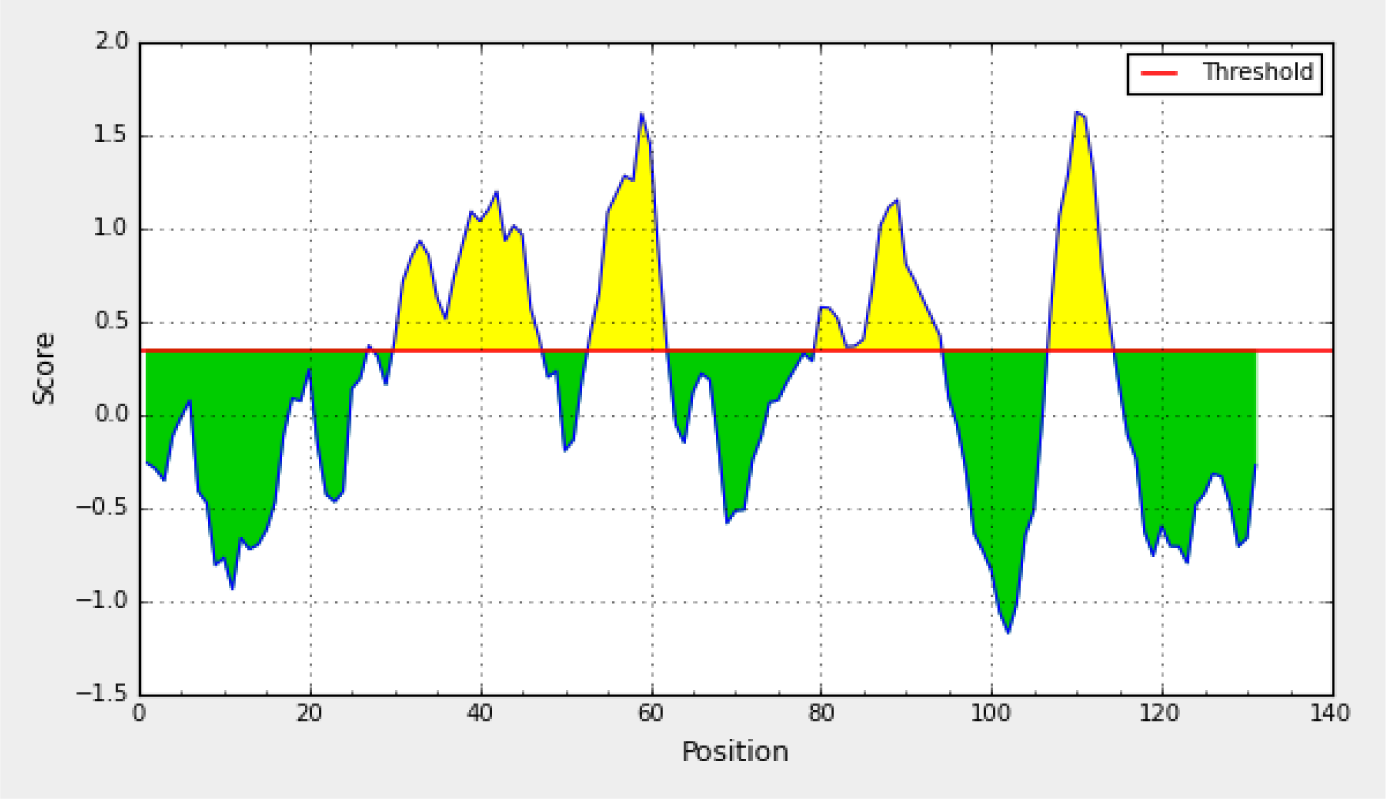
Potential B cell epitopes predicted from BepiPred tool. threshold value for for potential epitope was set 0.350. Regions shaded with yellow color are predicted as potential epitopes. The maximum score predicted here is 1.630.

### 3.4 Mapping of the B cell epitopes in the modeled structure confirms their presence on the surface of profilin-4

The predicted 3-D structure of profilin-4 was visualized (Figure 5 A) using Swiss PDB viewer tool [7]. Ramachandran plot was generated using an online tool RAMPAGE to validate the predicted structure (Figure 5 B) shows the amino acid distribution in different regions of the plot. Predicted B cell epitopes of the profilin-4 protein are mapped on the predicted 3D structure of the profilin-4 protein (Figure 6). The different coloured balls on the surface of the protein other than pink represent the 4 predicted B cell epitopes and regions in pink represents the core of the protein.

**Figure 5:**
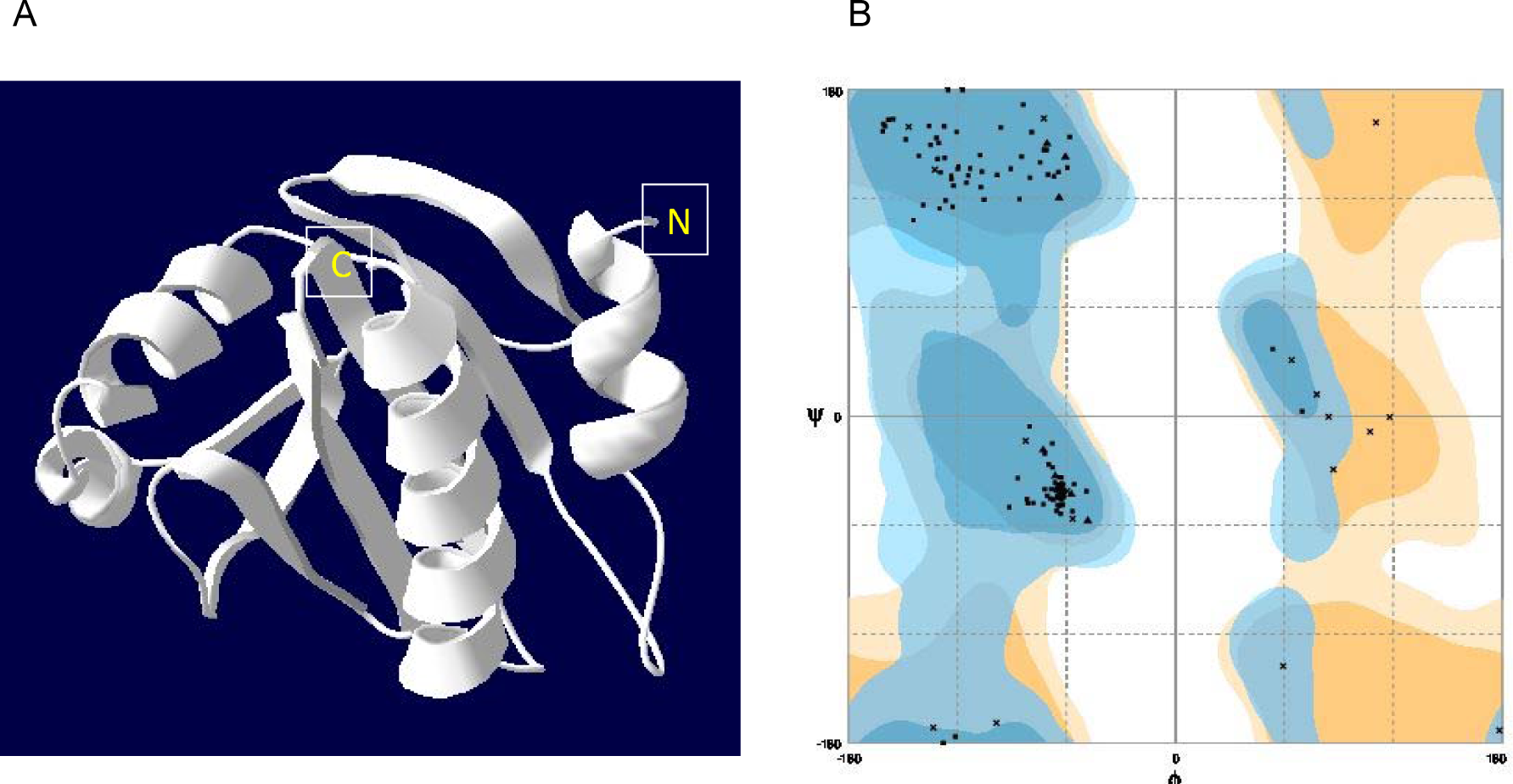
3D structure and its validation using Ramachandran plot for profilin-4 protein. (A) Cartoon representation of the predicted structure of the profilin-4 protein. This image has been developed using Swiss PDB viewer tool (B) amino acid residues are distributed in Ramachandran plot.

**Figure 6:**
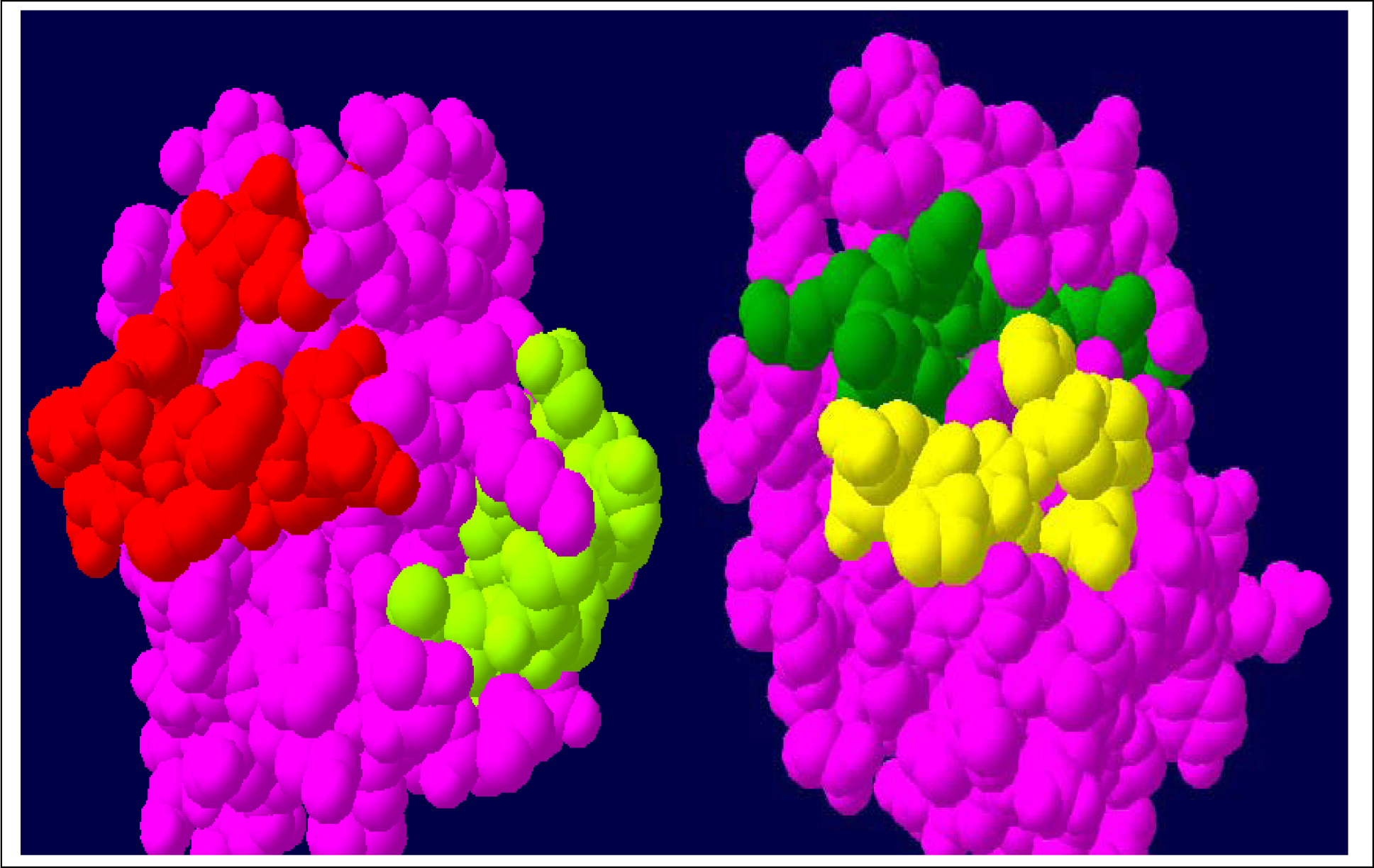
Mapping of B cell epitopes on the 3D structure of the profilin-4. BepiPred predicted epitopes are mapped on the surface of the profilin-4 protein structure where B cell epitopes are highlighted in red (epitope-1), olive (epitope-2), green (epitope-3) and yellow (epitope-4) and the rest of the non-reactive portion highlighted in pink.

## 4. Conclusion

Due to the increasing trend of maize production in the world (Figure 7), it’s urgent to analyze maize profilin-4 isoform’s potency as an allergen. In this study, it is evident that profilin-4 is a potential antigen, however, further in vitro analysis is required to validate along with the knowledge of the array of allergens in the pollen.

**Figure 7:**
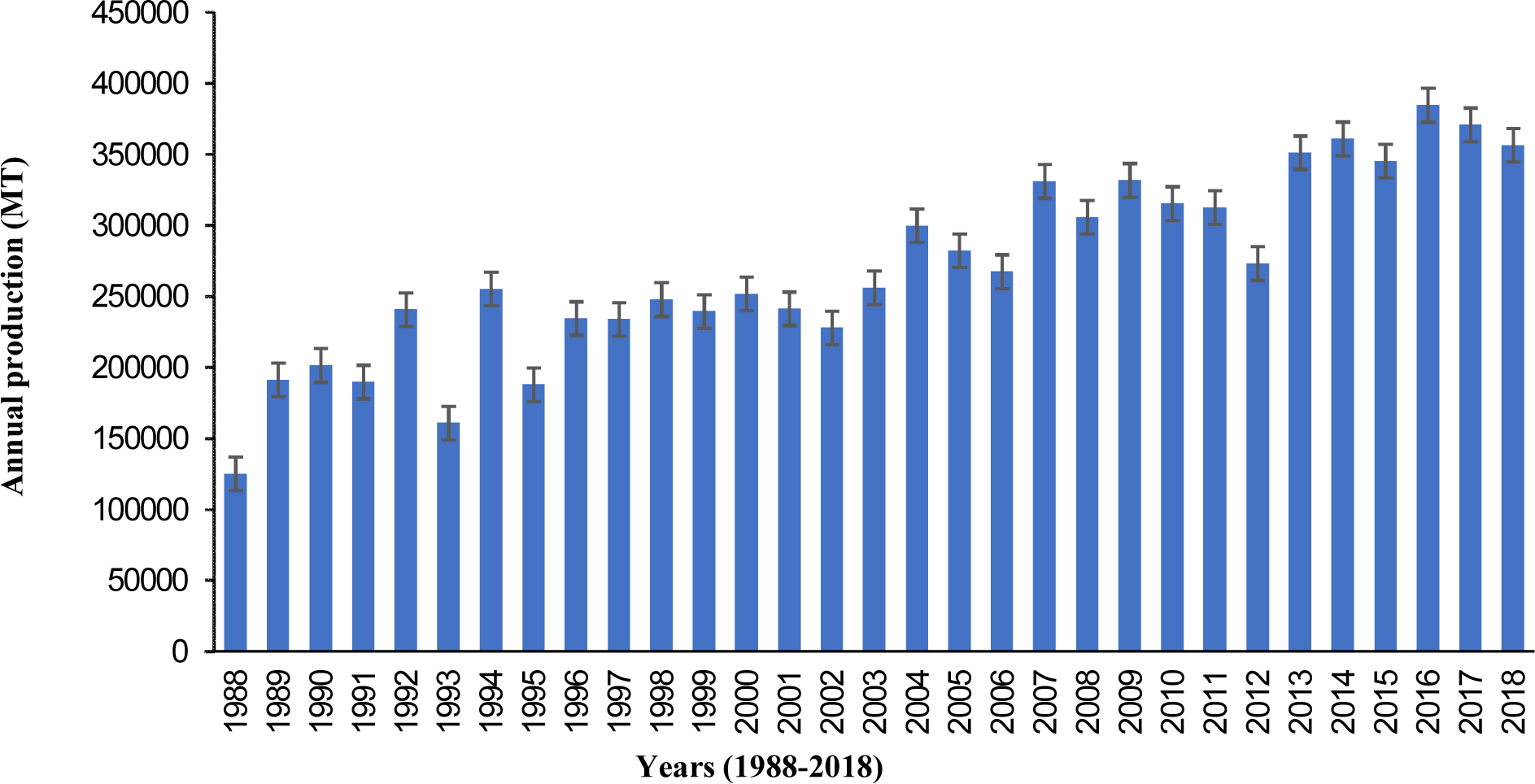
World corn production by from 1988 to 2018. This bar diagram shows a gradual increase of corn production over three decades around the world. (Modified from https://www.indexmundi.com/agriculture/?country=us&commodity=corn&graph=production)

## Acknowledgements

The authors are grateful to the Department of Biochemistry and Molecular Biology, University of Dhaka, Bangladesh and the Department of Biological Sciences, St. John’s University, Queens, New York 11439.

## References

[1]. S. B. Lehrer and J. E. Salvaggio, “Allergens: Standardization and Impact of Biotechnology—A Review,” Allergy Asthma Proc., vol. 11, no. 5, pp. 197–208, Sep. 1990.

[2]. C. Ozdemir, M. Akdis, and C. A. Akdis, “T-Cell Response to Allergens,” in Anaphylaxis, vol. 95, Basel: KARGER, 2010, pp. 22–44.

[3]. A. Vojdani, “Detection of IgE, IgG, IgA and IgM antibodies against raw and processed food antigens,” Nutr. Metab. (Lond)., vol. 6, no. 1, p. 22, May 2009.

[4]. J. A. Woodfolk, “T-cell responses to allergens,” J. Allergy Clin. Immunol., vol. 119, no. 2, pp. 280–294, Feb. 2007.

[5]. D. A. O. Taketomi, Ernesto A., Almeida, Karine C., Pereira, Fernando L., Silva, “Allergens: sources, exposure and sensitization levels, diagnostic tools and immunotherapeutical applications. J. Med. Med. Sci. 2.”

[6]. “FAOSTAT.” [Online]. Available: http://www.fao.org/faostat/en/#home. [Accessed: 05-Jan-2018].

[7]. H. Behrendt et al., “Air pollution and allergy: experimental studies on modulation of allergen release from pollen by air pollutants.,” Int. Arch. Allergy Immunol., vol. 113, no. 1–3, pp. 69–74.

[8]. A. Bufe, “The biological function of allergens: relevant for the induction of allergic diseases?,” Int. Arch. Allergy Immunol., vol. 117, no. 4, pp. 215–9, Dec. 1998.

[9]. P. Vallier, S. Balland, R. Harf, R. Valenta, and P. Deviller, “Identification of profilin as an IgE-binding component in latex from Hevea brasiliensis: clinical implications.,” Clin. Exp. Allergy, vol. 25, no. 4, pp. 332–9, Apr. 1995.

[10]. R. Valenta et al., “Identification of profilin as a novel pollen allergen; IgE autoreactivity in sensitized individuals.,” Science, vol. 253, no. 5019, pp. 557–60, Aug. 1991.

[11]. R. Asero et al., “Detection of clinical markers of sensitization to profilin in patients allergic to plant-derived foods.,” J. Allergy Clin. Immunol., vol. 112, no. 2, pp. 427–32, Aug. 2003.

[12]. S. L. Taylor and S. L. Hefle, “Will genetically modified foods be allergenic?,” J. Allergy Clin. Immunol., vol. 107, no. 5, pp. 765–771, May 2001.

[13]. S. M. Gendel, “Sequence databases for assessing the potential allergenicity of proteins used in transgenic foods.,” Adv. Food Nutr. Res., vol. 42, pp. 63–92, 1998.

[14]. S. M. Gendel, “The use of amino acid sequence alignments to assess potential allergenicity of proteins used in genetically modified foods.,” Adv. Food Nutr. Res., vol. 42, pp. 45–62, 1998.

[15]. J. D. Astwood and R. L. Fuchs, “Allergenicity of foods derived from transgenic plants.,” Monogr. Allergy, vol. 32, pp. 105–20, 1996.

[16]. D. D. Metcalfe, J. D. Astwood, R. Townsend, H. A. Sampson, S. L. Taylor, and R. L. Fuchs, “Assessment of the allergenic potential of foods derived from genetically engineered crop plants.,” Crit. Rev. Food Sci. Nutr., vol. 36 Suppl, pp. S165-86, 1996.

[17]. E. Gasteiger et al., “Protein Identification and Analysis Tools on the ExPASy Server,” in The Proteomics Protocols Handbook, Totowa, NJ: Humana Press, 2005, pp. 571–607.

[18]. A. S. Kolaskar and P. C. Tongaonkar, “A semi-empirical method for prediction of antigenic determinants on protein antigens.,” FEBS Lett., vol. 276, no. 1–2, pp. 172–4, Dec. 1990.

[19]. J. M. Parker, D. Guo, and R. S. Hodges, “New hydrophilicity scale derived from high-performance liquid chromatography peptide retention data: correlation of predicted surface residues with antigenicity and X-ray-derived accessible sites.,” Biochemistry, vol. 25, no. 19, pp. 5425–32, Sep. 1986.

[20]. J. Larsen, O. Lund, and M. Nielsen, “Improved method for predicting linear B-cell epitopes.,” Immunome Res., vol. 2, no. 1, p. 2, Apr. 2006.

[21]. M. Källberg et al., “Template-based protein structure modeling using the RaptorX web server,” Nat. Protoc., vol. 7, no. 8, pp. 1511–1522, Jul. 2012.

[22]. J. Ma, J. Peng, S. Wang, and J. Xu, “A conditional neural fields model for protein threading,” Bioinformatics, vol. 28, no. 12, pp. i59–i66, Jun. 2012.

[23]. N. Guex and M. C. Peitsch, “SWISS-MODEL and the Swiss-Pdb Viewer: An environment for comparative protein modeling,” Electrophoresis, vol. 18, no. 15, pp. 2714–2723, Dec. 1997.

[24]. S. C. Lovell et al., “Structure validation by Cα geometry: [,ψ and Cβ deviation,” Proteins Struct. Funct. Bioinforma., vol. 50, no. 3, pp. 437–450, Jan. 2003.

[25]. S. Singh, B. Taneja, S. S. Salvi, and A. Agrawal, “Physical Properties of Intact Proteins May Predict Allergenicity or Lack Thereof,” PLoS One, vol. 4, no. 7, p. e6273, Jul. 2009.

[26]. K. Shakushiro, Y. Yamasaki, M. Nishikawa, and Y. Takakura, “Efficient scavenger receptor-mediated uptake and cross-presentation of negatively charged soluble antigens by dendritic cells.,” Immunology, vol. 112, no. 2, pp. 211–8, Jun. 2004.

[27]. A. Bachmair, D. Finley, and A. Varshavsky, “In vivo half-life of a protein is a function of its amino-terminal residue.,” Science, vol. 234, no. 4773, pp. 179–86, Oct. 1986.

[28]. J. Söllner and B. Mayer, “Machine learning approaches for prediction of linear B-cell epitopes on proteins,” J. Mol. Recognit., vol. 19, no. 3, pp. 200–208, May 2006.

[29]. Tajamul Rouf Shah, Kamlesh Prasad, Pradyuman Kumar & Fatih Yildiz. “Maize-A potential source of human nutrition and health: A review”, Cogent Food & Agriculture vol. 2, Iss. 1, 2016

